# The N-terminal Domain of cpTatC Protein Interacts with the Precursor Mature Domain in Chloroplast TAT Translocation

**DOI:** 10.1101/2024.11.24.625085

**Authors:** Thilini Maddethalawe, Krystina Hird, Carole Dabney-Smith

**Author notes:** Corresponding author: Department of Chemistry and Biochemistry, Miami University, 651 East High St., Oxford, OH. Graduate Program in Biochemistry and Molecular Biology, Michigan State University, Lansing, Michigan, United States of America (Krystina Hird).

## Abstract

The chloroplast Twin Arginine Transport (cpTAT) protein translocation pathway is one of the thylakoid membrane’s two protein transport pathways for getting proteins into the lumen. The cpTAT system distinguishes itself by transporting fully folded proteins across the thylakoid, using the sole energy source of the proton motive force (PMF). The cpTAT pathway is evolutionarily conserved with the TAT pathway found in many bacteria and archaea. Although small differences exist, TAT systems in different organisms share homologous protein composition and similar molecular mechanisms. The cpTAT system comprises cpTatC, Hcf106, and Tha4 (the prokaryotic homologs of these proteins are TatC, TatB, and TatA, respectively). (cp)TatC is one of the essential proteins in the (cp)TAT system, as it is present in the receptor complex. The amino acid sequence alignment of cpTatC and TatC protein in archaea and *E. coli* has shown a unique N-terminal amino acid extension of 70-100 amino acids in mature cpTatC that is not present in prokaryotic TatC. However, the role of the amino-terminal extension in cpTatC function is still unknown. We present crosslinking evidence that the amino-terminal extension directly interacts with the precursor mature domain during TAT protein translocation and may serve to prime the transporter with bound precursor in the absence of the PMF.

**Highlights:**

- The cpTatC protein in the cpTAT pathway has a unique amino-terminal extension not found in bacterial or archaeal TatC proteins.
- The N-terminal extension of cpTatC interacts with the precursor mature domain during TAT translocation.
- The crosslinking data indicate that different regions within the N-terminal extension exhibit varying intensities when binding to the precursor.

## 1. Introduction

Chloroplasts are structurally complex organelles, with three membranes and three aqueous compartments. The intermembrane space of a chloroplast is enclosed by the chloroplast outer envelope membrane and chloroplast inner envelope membrane. The thylakoid membrane, a key component, is embedded in the aqueous stroma of the chloroplast, which is enclosed by the double envelope membranes. The innermost compartment of the chloroplast is the aqueous lumen, which is enclosed by the thylakoid membranes. Chloroplasts require between 2000 to 3000 proteins for full metabolic functionality, of which more than 97% are encoded by nuclear genes (Cline & Dabney-Smith, 2008, Cline & McCaffery, 2007, Leister, 2003, New *et al*., 2018). A subset of those are found in the thylakoid membrane and lumen, contain ∼ 100 different proteins: about 50% of the thylakoid membrane proteins and all the thylakoid lumen-resident proteins are encoded by genes in the nucleus, synthesized in the cytoplasm as higher molecular weight precursors, and their import occurs post-translationally (Jarvis & Robinson, 2004, Keegstra & Cline, 1999, Peltier *et al*., 2002, Schubert *et al*., 2002). The precursor comprises an N-terminal amino acid extension, the transit peptide, which contains the organellar targeting information that facilitates their transport to their final destination (Schatz & Dobberstein, 1996, von Heijne, 1990). Thylakoid-destined proteins have a bipartite transit peptide, meaning two signal sequences that act sequentially to allow import into the storms, followed by cleavage and exposure of the second sequence, and subsequent routing to the thylakoid (Cline & Theg, 2007, Schatz & Dobberstein, 1996).

Two major membrane-bound protein transport pathways work in parallel to import soluble proteins on the thylakoid membrane: the chloroplast general Secretary pathway (cpSEC) and the chloroplast Twin Arginine Transport pathway (cpTAT) (Celedon & Cline, 2013, Cline & Dabney-Smith, 2008, New et al., 2018). The cpSEC pathway is responsible for transporting mostly unfolded proteins, utilizing adenosine triphosphate (ATP) as the primary energy source enhanced by the presence of the proton motive force (PMF) (Dalbey & Robinson, 1999, Hynds *et al*., 1998, Mori *et al*., 1999, Mori & Ito, 2001). In contrast, the cpTAT system transports primarily tightly folded proteins (Creighton *et al*., 1995, Musser & Theg, 2000), using proton motive force (PMF) as the sole energy source in an NTP-independent manner (Celedon & Cline, 2013, Cline & Theg, 2007, Auldridge *et al*., 2006, New et al., 2018). cpTAT system is a vital protein translocation system for the higher plants as it enables the translocation of proteins involved in the photosynthetic processes, such as the oxygen-evolving complex of PSII, PsbP (23 kD; OE23), and PsbQ (17 kD; OE17) (Ifuku *et al*., 2011, Yi *et al*., 2006), subunit T of PSII (Kapazoglou *et al*., 1995), and subunit N of PSI (Haldrup *et al*., 1999).

The cpTAT system comprises three transmembrane proteins that work together to form active translocase. In the thylakoid, these three proteins are cpTatC, Hcf106, and Tha4, and in *E. coli*, the corresponding orthologues are TatC, TatB, and TatA (Cline & Dabney-Smith, 2008, Frain *et al*., 2016, Ma *et al*., 2018, New et al., 2018, Patel *et al*., 2014). Tha4 and Hcf106 are structurally similar and at the amino acid level but have distinguished functions in the TAT system. Both are nuclear-encoded proteins containing a transit peptide that targets the protein to the stroma; mature Tha4 is 8.9 kD, while mature Hcf106 is 19 kD. Both proteins are anchored in the lipid bilayer by the thylakoid lumen proximal amino-terminal transmembrane domain (TMD), followed by glycine-proline hinge region, amphipathic helix (APH), and acidic carboxy-terminal domain (C-tail). Moreover, Hcf106 possesses an extended APH and C-tail compared to Tha4. cpTatC is also encoded by the nuclear gene as a preprotein with transit peptide, and the pea isoform of the cpTatC is 33.3 kD with 303 amino acid residues (Cline & McCaffery, 2007, Ma et al., 2018, New et al., 2018, Pal *et al*., 2013) Based on the solved crystal structures for TatC from the bacterium *Aquifex aeolicus*, cpTatC is predicted to have a glove-like shape with a concave pocket on one side of the protein as it sits in the membrane (Ramasamy *et al*., 2013, Rollauer *et al*., 2012). The mature form of cpTatC comprises six transmembrane domains with both N- and C-termini facing the stromal side of the thylakoid. Previous cpTatC mutation studies have revealed that the signal peptide of the cargo/precursor proteins binds preferentially to the stromal proximal portions of TMD1 and TMD2 (Gérard & Cline, 2006). Furthermore, it has been demonstrated that amino acid changes in TMD5 impact the assembly of the overall complex (Ma & Cline, 2013, New et al., 2018).

The TAT system operates cyclically, and the TAT transport can be divided into discrete steps such as precursor binding to a cpTatC/Hcf106/Tha4 receptor complex (1:1:1), additional Tha4/TatA assembly with the precursor-bound receptor, precursor translocation, and Tha4/TatA disassembly. In its resting state, the translocon is a heterotrimer of tightly bound cpTatC and Hcf106 with loosely associated Tha4, which serves as the initial point of homo-oligomerization of additional Tha4 to create the translocation pore (Aldridge *et al*., 2012, Auldridge et al., 2006, Dabney-Smith & Cline, 2009, Pal et al., 2013). Assembly of additional Tha4 is PMF dependent after which the translocation of cargo occurs, followed by the disassembling of the Tha4 from the receptor complex, thus resetting the receptor complex for subsequent rounds of transport (Mori et al., 1999, Mori & Cline, 2002).

The structural and mechanical similarities between the bacterial TAT and cpTat systems are widely acknowledged (Müller & Klösgen, 2005, New et al., 2018). Phylogenetic analysis has revealed that nearly all TatC homologs are within the size range of 240-310 amino acid residues (Yen *et al*., 2002). However, the plant cpTatC protein exhibits a much longer N-terminal soluble domain of the mature protein, than those predicted for bacterial and algal TatC proteins (**Figure 1**) (Mori *et al*., 2001, Yen et al., 2002). This study aimed to investigate the role of the N-terminal extension of cpTatC through cysteine (Cys) scanning mutagenesis along the amino-terminal extension of mature cpTatC in *Pisum sativum* (garden pea). We employed a disulfide exchange crosslinking method to gain insights into the function of the N-terminal extension of pea cpTatC. During this study, we tested 29 different Cys-substituted cpTatC variants (**Figure 1B**) with tOE17A137C-(GS)_3_ (labeled as tOE17A137C further) (Pal et al., 2013) using disulfide crosslinking.

**Figure 1:**
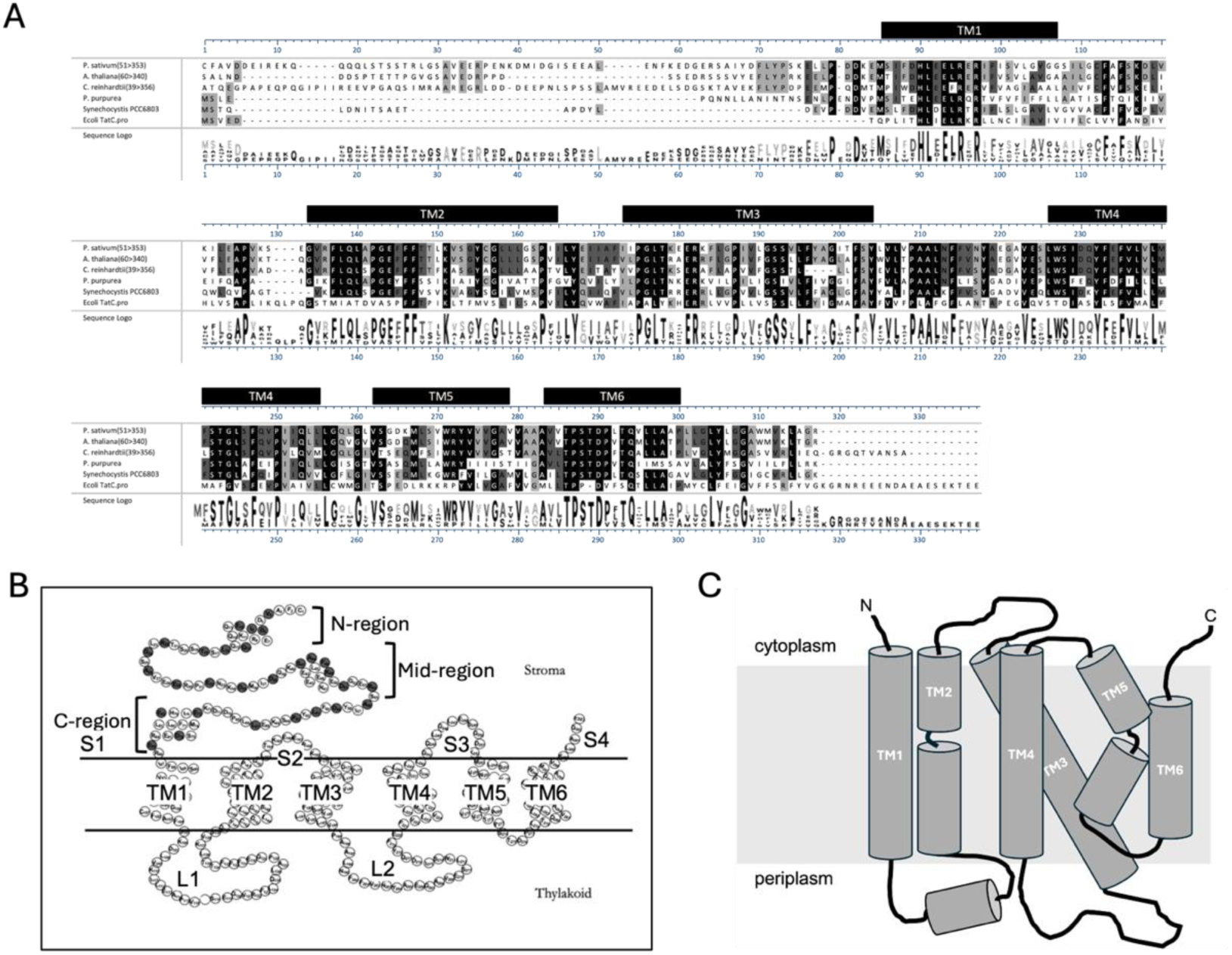
Membrane topology of cpTatC and sequence alignment of TatC from plant, alga, and bacteria. **A:** Amino acid sequence alignment of TatC homologs from *Pisum sativum (P. sativum)*, *Arabidopsis thaliana (A. thaliana), Chlamydomonas reinhardtii (C. reinhardtii), Porphyra purpurea (P. purpurea)*, *Synechocystis sp. PCC6803 (Synechocystis)*, and *Escherichia coli (E. coli)*. The alignment was generated in Lasergene DNAStar MegAlign Pro using MAFFT alignment tool and the BLOSUM80 scoring matrix. **B**: The filled circles indicate the positions of single cysteine substitutions on cpTaC generated in this study. The N-terminal extension of chloroplast TatC consists of amino acid residues 1-86 in the protein from *Pisum sativum.* Amino acid residues 1-30 are designated as the N-region; residues 31-60 are the mid-region; and residues 61-86 are the C-region. Residues ∼76-86 of the C-region are also considered stromal region 1 (S1), corresponding to the region in prokaryotic TatC proteins that make the N terminus. TatC proteins have six transmembrane domains (TM1-TM6) connected by loops extending into the stroma (S1-S4) and the lumen (L1-L2) of thylakoid. **C**: For comparison, the membrane topology of bacterial TatC with short N-terminal domain is shown (adapted from Patel et al., 2014).

## 2. Materials and Methods

### 2.1 Isolation of chloroplasts

Intact chloroplasts were isolated from 10-12-day-old pea (*Pisum sativum*) seedlings (Cline, 1986). Briefly, the isolated chloroplasts were suspended at 1 mg chlorophyll/mL with import buffer (IB; 50 mM HEPE-KOH, pH 8.0, 330 mM sorbitol) and kept on ice until used.

### 2.2 Generation of cpTatC and precursor Cys variants

Cys substituted cpTatC proteins (cpTatCXnC, where X is the relevant amino acid at the position of n has been substituted with the Cys) were generated by primer-based mutagenesis (New England Biolabs). The full-length precursor to cpTatC with the transit peptide from the precursor to the small subunit of RuBisCO in pGEM-4Z was used as the template (Aldridge et al., 2012, Aldridge *et al*., 2014). The three intrinsic cysteines (Cys) (C1, C101, and C142) were substituted with alanine (Ala) to generate cpTatC proteins with single Cys substitutions. cpTatC amino acid substitutions were verified by the Sanger sequencing (GENEWIZ). tOE17A137C precursor protein was described previously (Pal et al., 2013).

### 2.3 In-vitro translation of cpTatC Cys variants and precursor protein

cpTatC single Cys-substituted variants and the single Cys-substituted tOE17A137C GS3 precursor were radiolabeled with [^3^H] Leucine (Leu). All radiolabeled and unlabeled proteins (single Cys-substituted cpTatC variants and single Cys-substituted tOE17A137C precursors were *in vitro* translated by wheat germ extract (Promega) system with capped mRNA (Auldridge et al., 2006). The translated products were diluted with an equal volume of 30 mM Leu in 1X IB and 60 mM Leu in 2X IB.

### 2.3 DTNB derivatization of precursor

*In-vitro* translated radiolabeled and unlabeled tOE17A137C precursor was derivatized with 5, 5’-dithiobis (2-nitrobenzoic acid) (DTNB). The translated product of the precursor was incubated with 80 μM DTNB) at room temperature for 30 minutes before use in crosslinking (Pal et al., 2013, Tokatlidis *et al*., 1996).

### 2.4 cpTatC import assay

*In-vitro* translated radiolabeled and unlabeled cpTatC Cys-substituted variants were combined with 1 mg/mL chloroplast, 35 mM MgCl_2_, and 100 mM DTT. The import reaction was carried out in a 25°C water bath under 100 μE/m2/s of white light for 40 minutes. After the import, intact chloroplasts were isolated by centrifugation through 35% Percoll cushion and washed with 1X IB. Then, the thylakoid membrane was obtained by the osmotic lysis of intact chloroplast with HKM (50 mM HEPES-KOH, pH 8.0, 10 mM MgCl_2_) for 10 min on ice. After the incubation, 2X IBM (2X IB, ten mM MgCl_2_) was added, and thylakoids were pelleted at 3200 g for 8 minutes. Thylakoids were resuspended in 1X IBM at 1 mg/mL and used for the crosslinking assay (Mori & Cline, 1998).

### 2.5 Disulfide Crosslinking assay

cpTatC-integrated thylakoids were used for the crosslinking assays. Half of the radiolabeled cpTatC reaction was combined with the unlabeled tOE17A137C precursor, and the other half was combined with the 1X IBM as the no precursor control. The unlabeled cpTatC reaction was combined with the radiolabeled tOE17A137C precursor. Precursor and thylakoids were mixed in a 1:1 ratio and incubated in the dark at 4 °C for 30 minutes. Then the thylakoids were centrifuged at 3200 g for 8 minutes and resuspended in 1X IBM. 1 mM copper (II)-1, 10-phenanthroline (CuPh) was added as an oxidant to catalyze disulfide formation between proximal cysteine residues and incubated for 5 in the dark. The crosslinking reaction was stopped by adding 1 M N-ethyl maleimide (NEM) and centrifuging at 3200 g for 8 min. The samples were quenched by adding five mM EDTA and ten mM NEM in 1X IBM and centrifuged at 3200 g for 8 min. The thylakoid pellet was resuspended to 0.3 mg/mL chlorophyll and divided into two for analysis.

### 2.6 SDS-PAGE and Fluorography

2X non-reducing sample solubilization buffer (2X SSB-NR; 100 mM Tris-HCl (pH 6.8), 8 M urea, 5% SDS, and 30% glycerol, 0.1% bromophenol blue) was added to one-half of the samples and the second half was mixed with 2X reducing SSB (2X SSB-R; 100 mM Tris-HCl (pH 6.8), 8 M urea, 5% SDS, and 30% glycerol, 10% beta-mercaptoethanol, 0.1% bromophenol blue). Samples were analyzed using SDS-PAGE followed by fluorography.

## 3. Results

### 3.1 cpTatC single Cys-substituted variants import into the thylakoid membrane

To study the interaction between the N-terminal extension of cpTatC protein in land plants and the precursor mature domain during chloroplast TAT translocation, we generated a series of single Cys-substituted cpTatC variants (**Figure 1B**). We tested their import into the thylakoid membrane (**Figure 2**) to confirm the substitutions did not affect proper localization. All the single Cys-substituted variants successfully integrated into the thylakoid membrane. cpTatC E29C contained an internal truncation and was not used. **Figure 2** shows import and thylakoid localization for the five variants cpTatC E10C, R22C, R54C, V4C, and F48C. Precursors to cpTatC and Cys-substituted variants (∼37 kD in size; lanes 1-4, 6-7, 9-10, 12-15, 17-18, and 20), were imported into intact chloroplasts, processed to mature (∼28 kD) and localized to the thylakoid membrane (lanes 9, 12, 17, 18, and 20). The stromal fraction shows no cpTatC band, confirming successful integration into the thylakoid membrane (Lanes 5, 8, 11, 16, and 19). The wild type (WT) cpTatCaaa, translated *in-vitro*, can integrate into the thylakoid membrane (lane 1 of **supplementary figure 1**) when imported into intact chloroplasts (Aldridge et al., 2014). This shows similar integration behavior to the single Cys-substituted cpTatCaaa variants. The successful import of single Cys-substituted cpTatC variants into intact chloroplasts and thylakoid localization strongly suggests that the single cysteine substitutions do not have an adverse effect.

**Figure 2:**
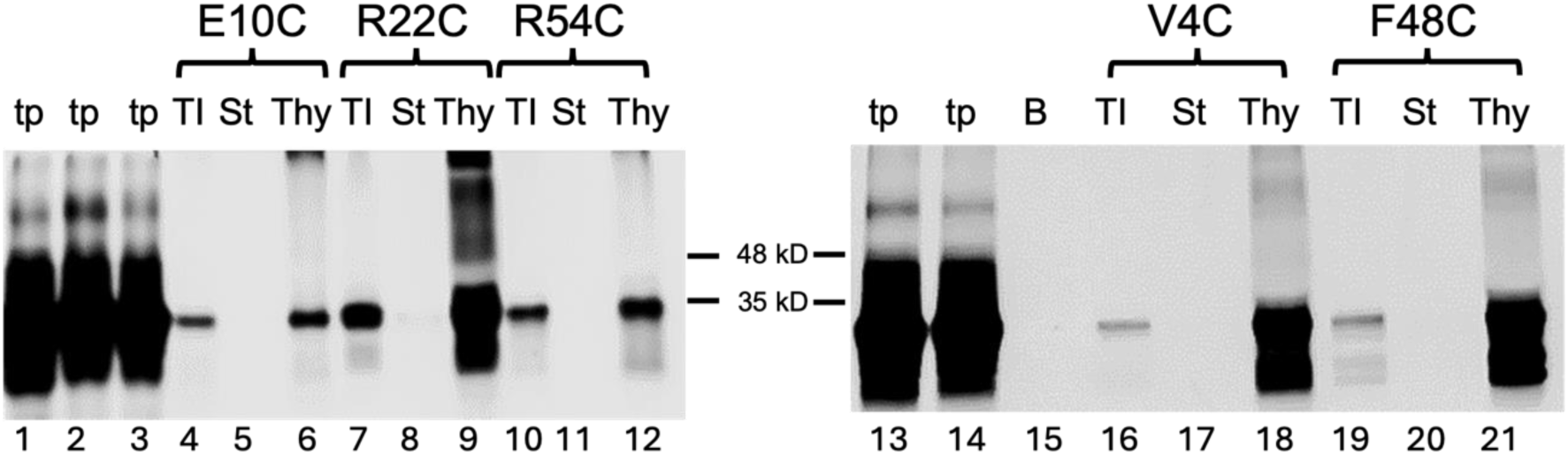
cpTatC single Cys substituted variants imported into the isolated chloroplast and integrated into the thylakoid membrane successfully. *In-vitro* translated pea cpTatC single Cys substituted variants were imported into the chloroplast and lysed to isolate the thylakoid membrane to see thylakoid integration of cpTatC single Cys substitutions. Lanes 1, 2, and 3 in the left panel are in-vitro translated products (tp) cpTatC E10C, R22C, and R54C, respectively. Lanes 13 and 14 of the right panel are tp of cpTatC V4C and cpTatC F48C, respectively. The chloroplast import/total import (TI) of cpTatC single Cys substitutions are shown in lanes 4, 7, and 10 on the left and lanes 16 and 19 on the right panel. Each variant’s stromal fraction (St) shows on lanes 5, 8, and 11 on the left and lanes 17 and 20 on the right. Lanes 6, 9, and 12 in the left panel and lanes 18 and 21 in the right are each variant’s thylakoid fraction (Thy).

### 3.2 Amino-proximal portion of N-terminal extension interacts with precursor during cpTAT transport

The thylakoid integration assays confirm that the thylakoid import ability of the protein remains uncompromised following single Cys substitutions. After validating the functionality of each single Cys-substituted cpTatC, Cys disulfide crosslinking assays are conducted accordingly. Despite the successful integration and localization of the WT cpTatCaaa into the thylakoid membrane, it does not exhibit interaction with the single Cys-substituted precursor, tOE17A137C-(GS)3 (**Supplementary** Figure 1).

To study the interaction between N-terminal domain of cpTatC, a single Cys-substituted precursor, tOE17A137C-(GS)_3_, was used. The tOE17 is the precursor of the 17kD subunit of the oxygen-evolving complex (OE17) with a truncated signal peptide lacking the acidic amino-proximal extension (Henry *et al*., 1997, Ma & Cline, 2000) and with a cysteine substitution at position 137 and containing a (GSSS)_3_ motif at the carboxyl terminus. The thiols on precursors were modified with 5,5’-dithiobis(2-nitrobenzoic acid) (DTNB) to prevent the formation of disulfide bonds between precursor proteins, while still allowing reaction with unmodified thiols in neighboring proteins (Pal et al., 2013, Tokatlidis et al., 1996). Pal et al. (2013) showed that the DTNB derivatized single Cys-substituted precursor, tOE17A137C-(GS)_3_, is competent for transport across thylakoid.

We employed Cys-Cys disulfide crosslinking assays with each variant to test the interaction between the N-terminal extension of cpTatC and the precursor mature domain, tOE17A137C. Precursor to cpTatC or the cysteine variants was expressed in-vitro as either radiolabeled (^3^H) or unlabeled versions. Labeled or unlabeled precursors to cpTatC or variants were imported into intact chloroplast in the illuminated water bath and the thylakoids harvested after import by hypotonic lysis (Aldridge et al., 2014, Celedon & Cline, 2012). The tOE17A137C precursor was derivatized with DTNB in order to prevent crosslinking with other tOE17A137C precursors (Pal et al., 2013). Harvested thylakoid were used to follow transport and crosslinking of either labeled or unlabeled tOE17A137C to labeled (**Figure 3**, lanes 2, 4-5, 7-8, left panel) or unlabeled cpTatC variants (lanes 3, 6, and 9, left panel). The radiolabeled cpTatC variants were combined with unlabeled precursors (**Figure 3**, lanes 4, 7, and 10, left panel). The use of CuPh oxidant improves the efficacy of crosslinking bond formation involving Cys residues located within approximately 5 Å of each other. This leads to a detectable shift in mobility on SDS-PAGE. We assessed cysteines placed in three regions of the N-terminal extension: the N-region, mid-region, and C-region (**Figure 1B**). The crosslink between cpTatC D6C-tOE17A137C (**Figure 3**, lanes 3 and 4, left panel) and cpTatC A26C-tOE17A137C (**Figure 3**, lanes 6 and 7, left panel) was evident by the characterized band shift at M_r_ ∼48kD, which correlates to an adduct of mature cpTatC (M_r_ ∼28 kD) and pre-tOE17A137C (M_r_ ∼20kD). cpTatC dimers (∼60 kD) appeared only in the samples containing radiolabeled cpTatC variants (**Figure 3**, lanes 4, 7, and 9, left panel).

**Figure 3:**
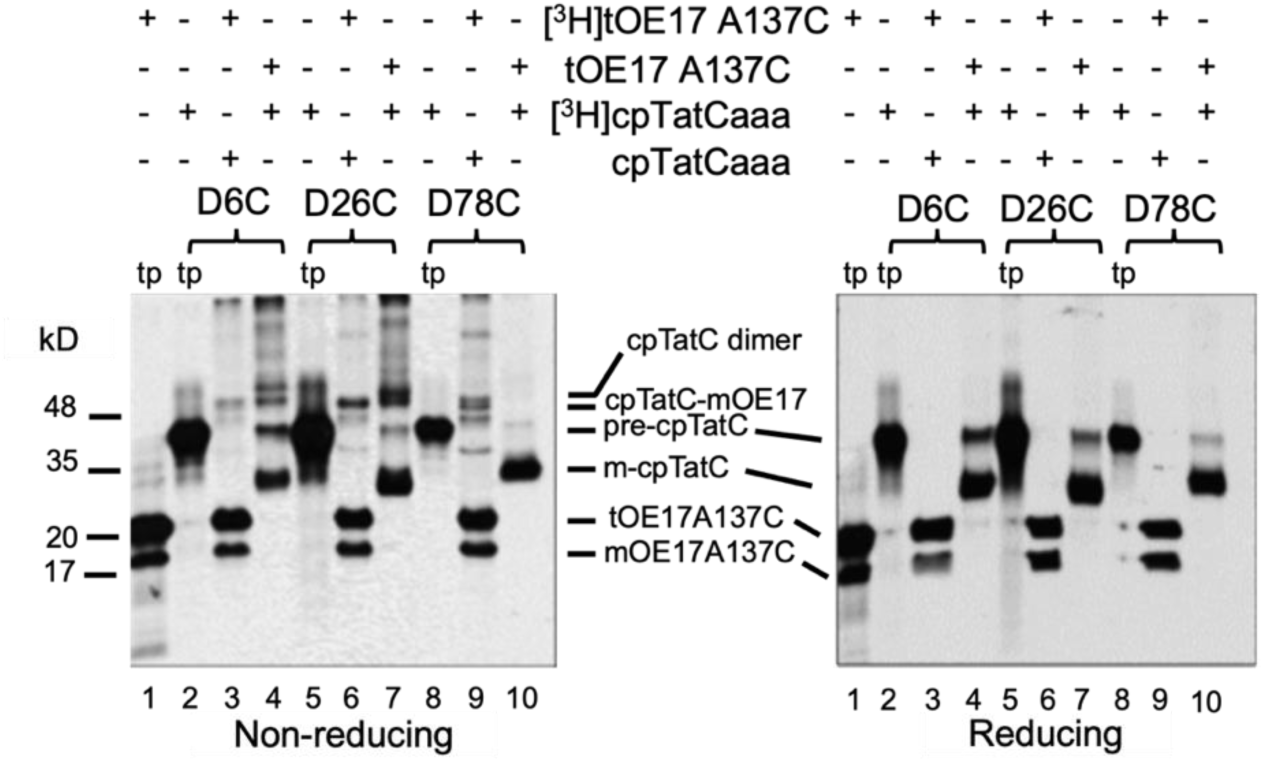
Single Cys substitute cpTatC D6C and A26C crosslinked with the tOE17A137C precursor. Lane 1 in the left and right panels is the translated product (tp) of the tOE17A137C[3H] precursor used in the disulfide crosslinking assays. Lanes 2, 5, and 8 in the left and right panels are translated products (tp) of the cpTatC single Cys substituted variants, as stated at the top of the figures. Isolated thylakoid containing radiolabeled single Cys substituted cpTatC variants were combined with unlabeled tOE17A137 (lanes 4, 7, and 10 in both panels). In contrast, isolated thylakoids containing unlabeled single Cys substituted cpTatC variants were combined with radiolabeled tOE17A137 (lanes 3, 6, and 9 in both panels). Precursor cpTatC (pre-cpTatC) shows in the gel at ∼37 kD, and mature cpTatC (m-cpTatC) at ∼28 kD. Precursor tOE17A136C (pOE17A137) shows at ∼20 kD, and mature tOE17A137C (mtOE17A137C) shows at 17 kD. Crosslinking assays were carried out as described in the Materials and Methods section and analyzed by 12.5% SDS-PAGE under non-reducing conditions (left panel) and reducing conditions (right panel).

The band shift results for cpTatC A26C-precursor (**Figure 3**, lanes 6 and 7, left panel) show higher band intensity than the gel band shift relevant for cpTatC D6C-precursor (**Figure 3**, lanes 3 and 4, left panel). It can be concluded that cpTatC A26C strongly interacts with the precursor during chloroplast TAT translocation compared to cpTatC D6C. When the samples were treated with β-mercaptoethanol-containing sample solubilization buffer (SSB), disulfide reduction caused the disappearance of all crosslinks, including cpTatC-tOE17A137C crosslinks as well as cpTatC dimers (**Figure 3**, lanes 3, 4, 6, and 7, right panel). The cpTatC D78C does not interact with the precursor (**Figure 3**, lanes 9 and 10 in the left panel).

Additionally, it was observed that cpTatC T18C, R22C, and E32C directly interact with the tOE17A137C during cpTat transport (**Supplementary** Figure 2). Based on the gel shift band intensity, it is evident that cpTatC R22C demonstrates a more pronounced interaction with the precursor (lanes 7 and 8 of the left panel in **Supplementary** Figure 2) as compared to cpTatC T18C (lanes 3 and 4 of the left panel in **Supplementary** Figure 2) and E32C (lanes 11 and 12 of the left panel in **Supplementary** Figure 2).

### 3.3 The middle region of the N-terminal extension interacts with precursor during cpTAT transport

Moving along the sequence of the N-terminal extension, we tested the interaction between the mid-region and precursor. Thylakoid integration and crosslinking assays with cpTatC S41C and K49C were conducted as described above. The band shift at M_r_ ∼48 kD (**Figure 4**, lanes 3 and 4, left panel) confirms the interaction between cpTatC S41C with tOE17A137C, which disappeared in the disulfide reduction condition (**Figure 4**, lanes 3 and 4, right panel). The intensity of the interaction between the cpTatC S41C-precursor (**Figure 4**, lanes 3 and 4, left panel) is similar to the intensity of the cpTatC A26C-precursor (**Figure 3**, lanes 6 and 7-9, left panel). The cpTatC K49C variant does not interact with the precursor (**Figure 4**, lanes 6 and 7, left panel).

**Figure 4:**
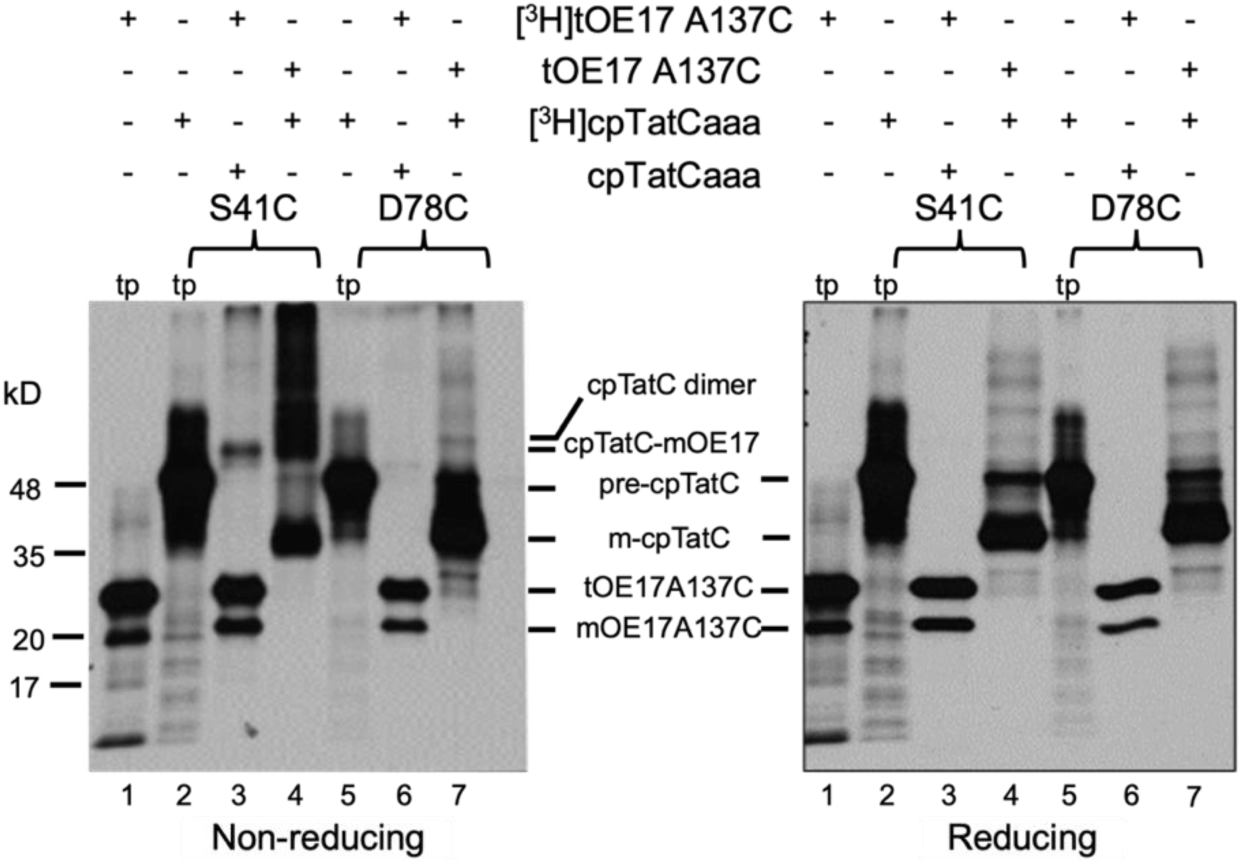
Single Cys substitute cpTatC S41C crosslinked with the tOE17A137C precursor. Lane 1 in the left and right panels is the translated product (tp) of the tOE17A137C[^3^H] precursor used in the disulfide crosslinking assays. Lanes 2 and 5 in the left and right panels are translated products (tp) of the cpTatC single Cys substituted variants, as stated at the top of the figures. Isolated thylakoid containing radiolabeled single Cys substituted cpTatC variants were combined with unlabeled tOE17A137(lanes 4 and 7 in both panels). In contrast, isolated thylakoids containing unlabeled single Cys substituted cpTatC variants were combined with radiolabeled tOE17A137 (lanes 3 and 6 in both panels). Precursor cpTatC (pre-cpTatC) shows in the gel at ∼37 kD, and mature cpTatC (m-cpTatC) at ∼33 kD. Precursor tOE17A136C (pOE17A137) shows at ∼ 25 kD, and mature tOE17A137C (mtOE17A137C) shows at 17 kD. Crosslinking assays were carried out as described in the Materials and Methods section and analyzed by 12.5% SDS-PAGE under non-reducing conditions (left panel) and reducing conditions (right panel).

In addition, we conducted testing on the cpTatC D35C variant within the medium region of the N-terminal extension (**Supplementary** Figure 3). Band shifts at approximately 48 kD for the cpTatC D35C variant (lanes 3 and 4 of the **supplementary figure 3**) indicate a comparable interaction strength with the cpTatC R22C-precursor (lanes 7 and 8 of the left panel in **Supplementary** Figure 2), reflecting a medium-level strength.

In continuation, we have conducted further analysis of the Cys disulfide crosslinking assays involving cpTatC A44C, D51C, R54C, and L61C within the middle region. Upon evaluating the gel band shift intensity, we have determined that both cpTatC A44C (lanes 7 and 8 of the left panel of **supplementary figure 4**) and D51C (lanes 11 and 12 of the left panel of **supplementary figure 4**) demonstrate weak binding with the tOE17A137C precursor during chloroplast TAT transport. Additionally, our findings indicate that cpTatC R54C (lanes 3 and 4 of the left panel of **supplementary figure 5**) and L61C (lanes 7 and 8 of the left panel of **supplementary figure 5**) also exhibit weak binding with the tOE17A137C precursor.

### 3.4 cpTatC Q12C and L16C interact with the tOE17A137C during chloroplast TAT translocation

The crosslinking assay between cpTatC Q12C, L16C, and tOE17A137C shows that both Q12C and L16C variants crosslink with the precursor during chloroplast TAT translocation (lanes 3, 4, 7, and 8 in the left panel of **Figure 5**). Band shifts at approximately 48 kD for both variants indicate a similar interaction strength between cpTatC Q12C-precursor and cpTatC L16C-precursor.

**Figure 5:**
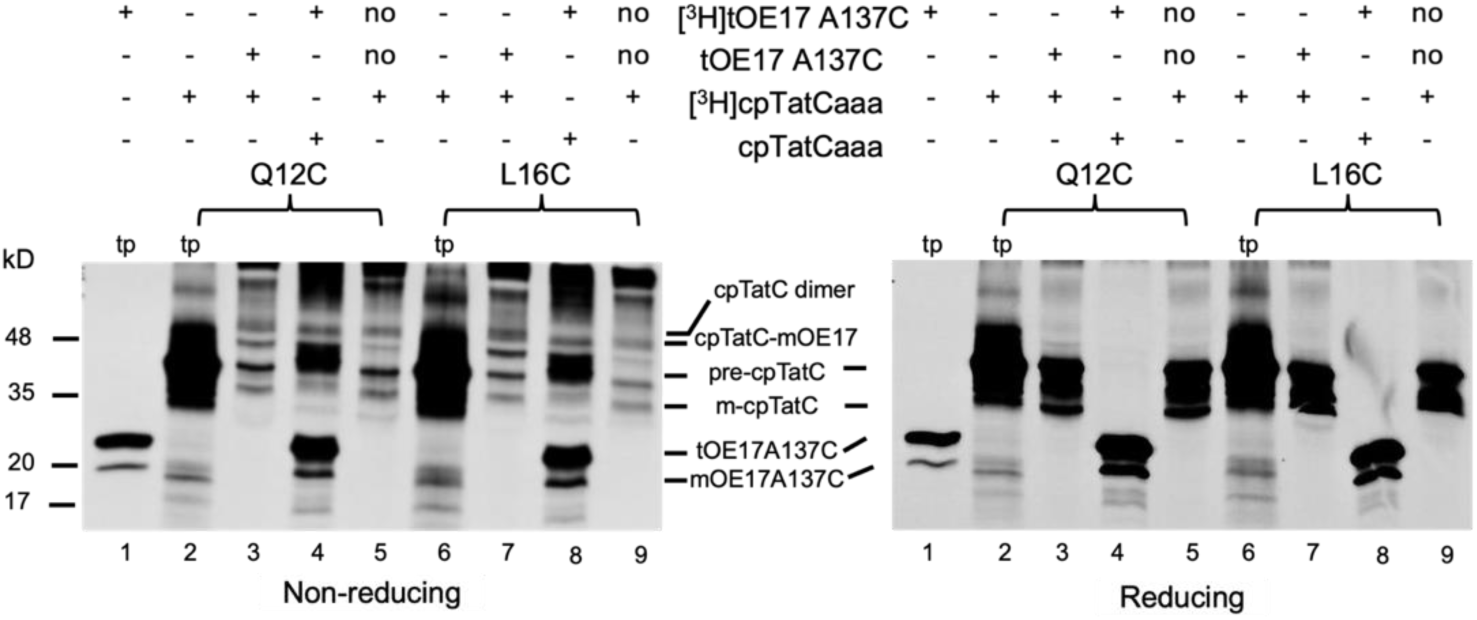
Single Cys substitute cpTatC Q12C and L16C crosslinked with the tOE17A137C precursor. Lane 1 in the left and right panels is the translated product (tp) of the tOE17A137C[3H] precursor used in the disulfide crosslinking assays. Lanes 2 and 6 in the left and right panels are translated products (tp) of the cpTatC single Cys substituted variants, as stated at the top of the figures. Isolated thylakoid containing radiolabeled single Cys substituted cpTatC variants were combined with unlabeled tOE17A137(lanes 3 and 7 in both panels). In contrast, isolated thylakoids containing unlabeled single Cys substituted cpTatC variants were combined with radiolabeled tOE17A137 (lanes 4 and 8 in both panels). As a control, radiolabeled cpTatC variants were combined with the same volume of 1X IBM (no precursor control) (lanes 5 and 9 in both lanes). Precursor cpTatC (pre-cpTatC) shows in the gel at ∼37 kD, and mature cpTatC (m-cpTatC) at ∼33 kD. Precursor tOE17A136C (pOE17A137) shows at ∼ 25 kD, and mature tOE17A137C (mtOE17A137C) shows at 17 kD. Crosslinking assays were carried out as described in the Materials and Methods section and analyzed by 12.5% SDS-PAGE under non-reducing conditions (left panel) and reducing conditions (right panel).

### 3.5 cpTatC I8C and E10C interact with the tOE17A137C during chloroplast TAT translocation

Building on the previous findings, we further probe the crosslinking between the precursor with cpTatC I8C and E10C. The gel band relevant to the cpTatC E10C-precursor (lanes 7 and 8 in the left panel of **Figure 6**) shows higher intensity than the cpTatC I8C-tOE17A137C (lanes 3 and 4 in the left panel of **Figure 6**), indicating a potentially stronger interaction.

**Figure 6:**
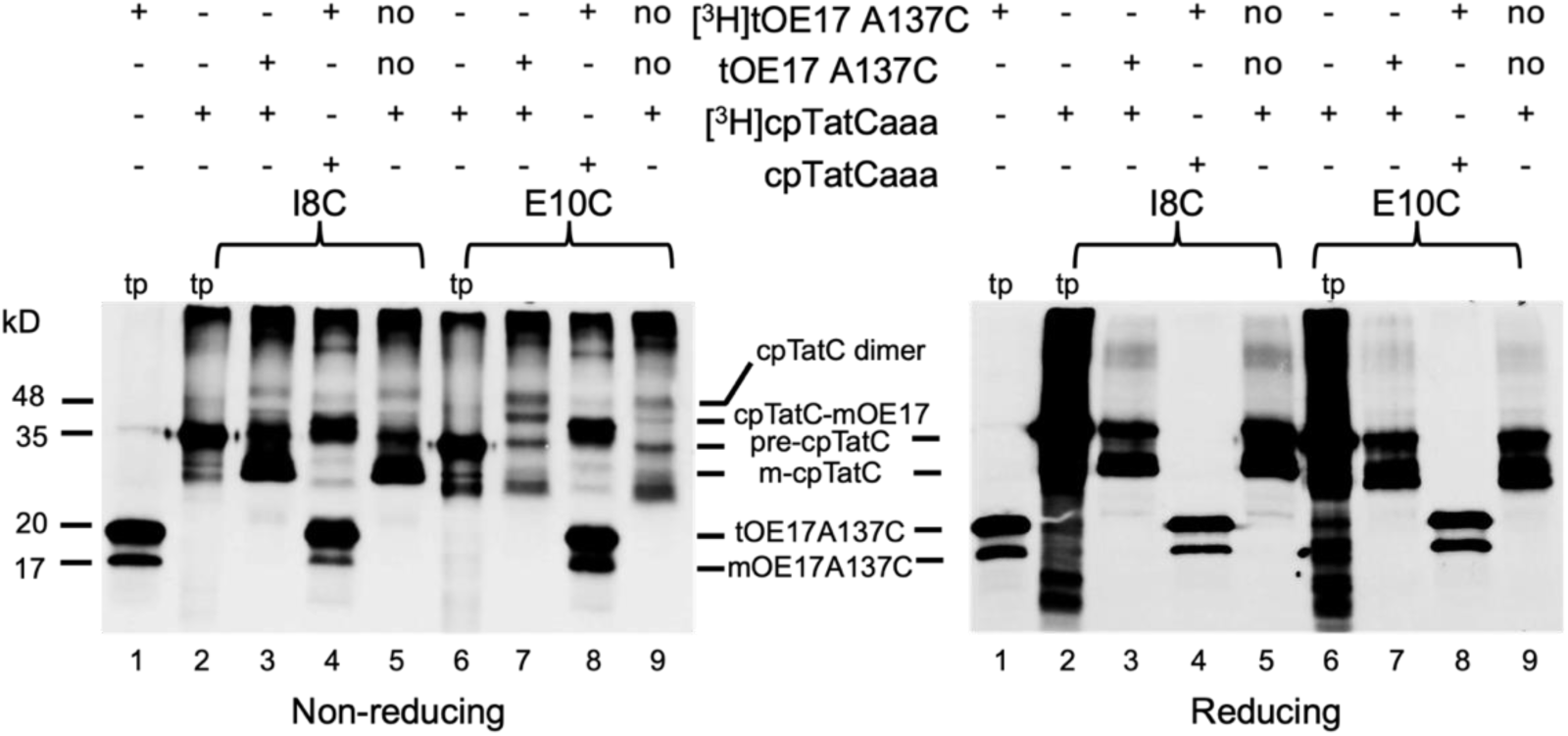
Single Cys substitute cpTatC I8C and E10C crosslinked with the tOE17A137C precursor. Lane 1 in the left and right panels is the translated product (tp) of the tOE17A137C[^3^H] precursor used in the disulfide crosslinking assays. Lanes 2 and 6 in the left and right panels are translated products (tp) of the cpTatC single Cys substituted variants, as stated at the top of the figures. Isolated thylakoid containing radiolabeled single Cys substituted cpTatC variants were combined with unlabeled tOE17A137C (lanes 3 and 7 in both panels). In contrast, isolated thylakoids containing unlabeled single Cys substituted cpTatC variants were combined with radiolabeled tOE17A137 (lanes 4 and 8 in both panels). As a control, radiolabeled cpTatC variants were combined with the same volume of 1X IBM (no precursor control) (lanes 5 and 9 in both lanes). Precursor cpTatC (pre-cpTatC) shows in the gel at ∼37 kD, and mature cpTatC (m-cpTatC) at ∼33 kD. Precursor tOE17A136C (pOE17A137) shows at ∼ 25 kD, and mature tOE17A137C (mtOE17A137C) shows at 17 kD. Crosslinking assays were carried out as described in the Materials and Methods section and analyzed by 12.5% SDS-PAGE under non-reducing conditions (left panel) and reducing conditions (right panel).

### 3.6 cpTatC G24C and V4C interact with the tOE17A137C during chloroplast TAT translocation

Figure 7 shows that the cpTatC G24C and V4C directly interact with the tOE17A137C during cpTat transport, but the cpTatC F48C variant does not interact with the precursor. Based on the gel band shift intensity, it can be concluded that cpTatC G24C binds strongly with the precursor while V4C interacts weakly with the tOE17A137C (Lanes 3, 4, 7, and 8 in the left panel of Figure 7).

**Figure 7:**
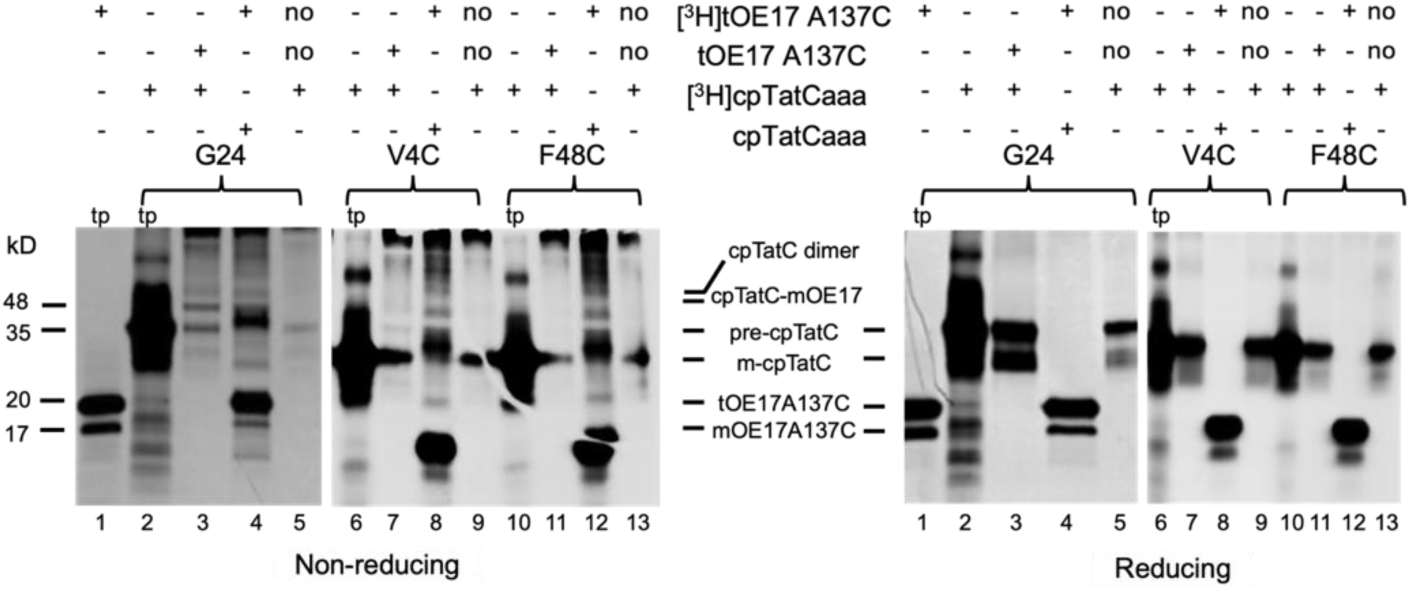
Single Cys substitute cpTatC E32C and V4C crosslinked with the tOE17A137C precursor. Lane 1 in the left and right panels is the translated product (tp) of the tOE17A137C[3H] precursor used in the disulfide crosslinking assays. Lanes 2, 6, and 10 in the left and right panels are translated products (tp) of the cpTatC single Cys substituted variants, as stated at the top of the figures. Isolated thylakoid containing radiolabeled single Cys substituted cpTatC variants were combined with unlabeled tOE17A137(lanes 3, 7, and 11 in both panels). In contrast, isolated thylakoids containing unlabeled single Cys substituted cpTatC variants were combined with radiolabeled tOE17A137 (lanes 4, 8, and 12 in both panels). As a control, radiolabeled cpTatC variants were combined with the same volume of 1X IBM (no precursor control) (lanes 5, 9, and 13 in both lanes). Precursor cpTatC (pre-cpTatC) shows in the gel at ∼37 kD, and mature cpTatC (m-cpTatC) at ∼33 kD. Precursor tOE17A136C (pOE17A137) shows at ∼ 25 kD, and mature tOE17A137C (mtOE17A137C) shows at 17 kD. Crosslinking assays were carried out as described in the Materials and Methods section and analyzed by 12.5% SDS-PAGE under non-reducing conditions (left panel) and reducing conditions (right panel).

## 4. Discussion

The mature domain of cpTatC possesses an N-terminal extension as compared to E. coli and algae TatC, the role of this extension in the chloroplast Tat system remains a mystery. In our study, we employed a crosslinking method, which allowed us to map interactions between the mature domain of the precursor (tOE17A137C) and the N-terminal extension of cpTatC through the formation of a disulfide bond between Cys within ∼5 Å of each other. We constructed a pool of single Cys-substituted cpTatC variants (Figure 1B) along the amino-terminal extension to study the direct interaction with the mature domain of a precursor during translocation. Studying the (cp) TAT system is a complex task, particularly due to the rapidity of the protein translocation process. To overcome the challenge of the rapid process, we employed a disulfide crosslinking assay. This technique can lock the formed protein complex by forming covalent crosslinks, facilitating our probing of this intricate process.

The (cp)TatC protein is a primary component of the (cp)TAT system. Multispanning cpTatC proteins bind tightly with Hcf106 and Tha4 (Aldridge *et al*., 2014) to create the receptor complex (Behrendt *et al*., 2007, Orriss *et al*., 2007), and it binds to the twin arginine motif (RR) of the signal peptide of the precursor (Alami *et al*., 2003, Gérard & Cline, 2006, Ma & Cline, 2013). Given that cpTatC is a multi-spanning membrane protein, it possesses the capacity to serve as a motor, thereby supplying the necessary force to aid in the movement of cargo through the translocation pore. Based on previous studies, it can be postulated that cpTatC provides a driving force to the movement of transmembrane protein through the translocon pore (Brüser & Sanders, 2003, Cline & McCaffery, 2007, Auldridge et al., 2006), which is composed of oligomers of Tha4 bound to the receptor complex. Oligomers of Tha4 assemble in response to binding of the precursor to the receptor complex and the presence of the proton motive force across the bioenergetic membrane. Mutation studies show that the RR motif of the signal peptide of the precursor binds specifically with the N-terminal stromal loop 1 (S1) and stromal loop 2 (S2) of the cpTatC protein in the receptor complex (Gérard & Cline, 2007, Ma & Cline, 2013). Previous studies showed the precursor binding simultaneously to both cpTatC and Hcf106, but signal peptide recognition is done by the S1 and S2 domains of the cpTatC initially (Ma & Cline, 2013).

The crosslinking data obtained from this study indicates that the N-terminal extension of the cpTatC protein interacts with the tOE17A137C precursor protein. The interaction involves specific cysteine residues, namely cpTatC V4C, D6C, I8C, E10C, Q12C, L16C, T18C, R22C, G24C, A26C, E32C, D35C, S41C, A44C, D51C, R54C, and L61C. Interestingly, different Cys residues of the amino-terminal extension interact with different binding intensities with the precursor. Based on the cpTatC-tOE17A137C crosslink band intensity, we classified the strength of the interaction (Figure 8). The S1 proximal residues (we tested cpTatC P63C, L67C, E73C, D78C, E82C, and E85C) of the cpTatC do not interact with the tOE17A137C at all. These data align with the findings of X. Ma & Cline, Ma and Cline (2013) which showed that signal peptide of the substrate variants tOE17-25C-20F or tOE17-20F-18C interacts with S1 proximal residues strongly, specifically with cpTatC L68C, E73C, D78C, and E85C. Based on the findings of this study, the interaction of the N-terminal extension of cpTatC with the precursor’s mature domain suggests that the N-terminal extension plays a role in stabilizing and positioning the precursor’s signal peptide with the stromal domains of the cpTatC.

**Figure 8:**
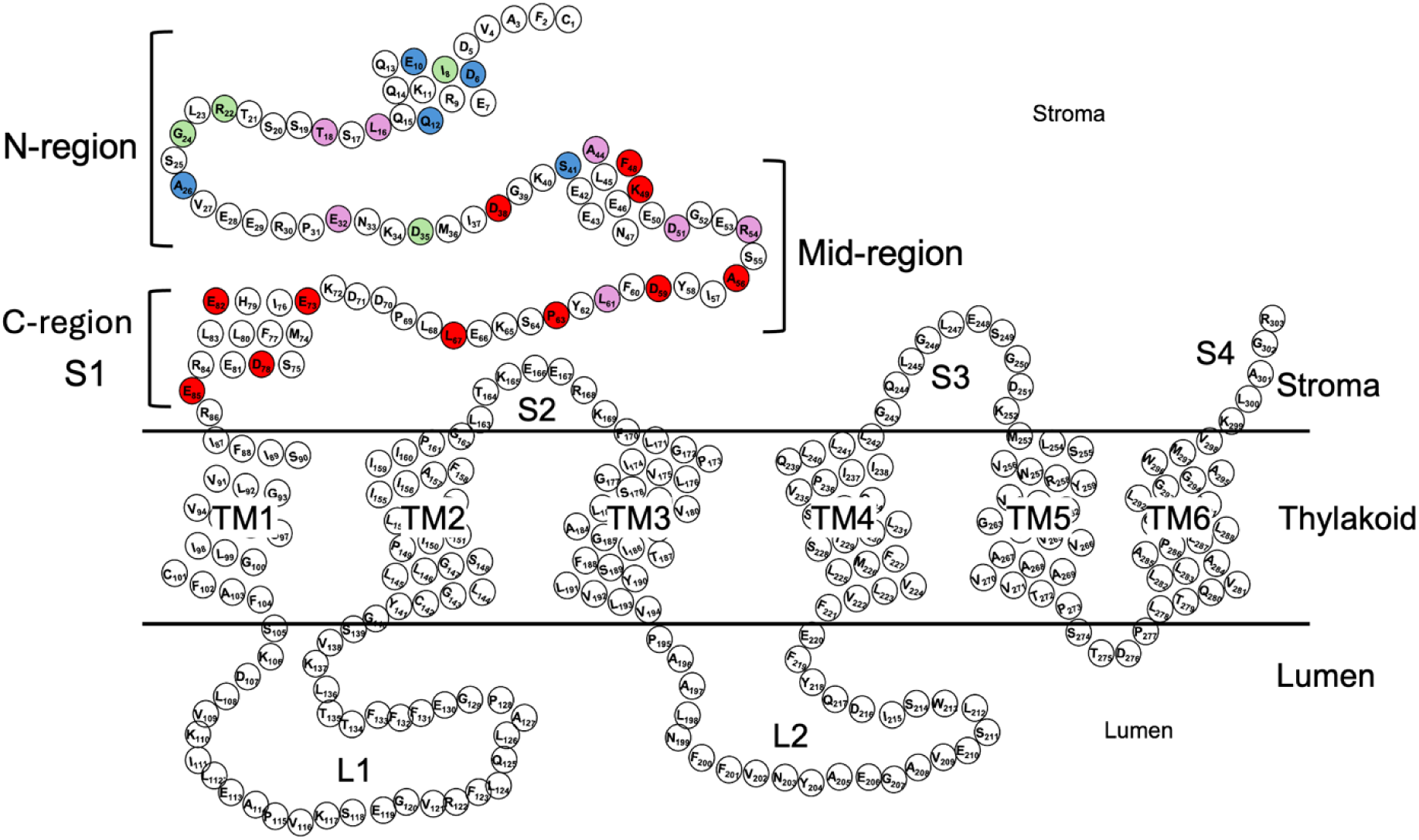
Heat map showing the strength of interaction between each cpTatC variant with tOE17A137C. Blue circles indicate a strong interaction between cpTatC and the precursor mature domain, green indicates an intermediate interaction, and pink indicates a weak interaction. Red circles indicate that the residue of cpTatC does not interact with the precursor. Labels on cpTatC are the same as those in Figure 1B.

The sum of currently available data revealed that the mechanism of the (cp)TAT system agrees strongly with the membrane weakening and the pulling mechanism (Brüser & Sanders, 2003) of the substrate through the TAT translocase. Firstly, the RR motif of the signal peptide binds with S1 and S2 of the cpTatC, which has been shown to be the primary recognition component of the RR consensus motif of the signal sequence (Gérard & Cline, 2006). In E. coli, it has also been shown that TatC directs the orientation of the RR-signal sequence, and TatC by itself shows a signal sequence insertase activity (Frobel *et al*., 2012). The same study suggested that TatC drives the hairpin-like embedment of RR-precursor protein through the translocon (Frobel et al., 2012). The cross-linking data collected throughout this study show that the precursor mature domain interacts with the N-terminal extension of the cpTatC throught the N-region and the mid-region but not the C-region (Figure 8).

Presumably, the N-terminal extension first binds with the precursor protein and that cpTatC-precursor binding stabilizes and possibly orients the precursor protein to facilitate the specific recognition and binding of the RR consensus motif of the signal peptide of the precursor protein to the S1 domain, which in turn initiates the formation of the hairpin loop of the signal peptide with cpTatC-Hcf106 receptor complex (Fincher *et al*., 1998, Frobel et al., 2012, Hou *et al*., 2006). The formation of the cpTatC-Hcf106-precursor complex triggers the formation of a translocation conduit by assembling Tha4 proteins when the thylakoid membrane is energized with PMF (Aldridge et al., 2014, Auldridge et al., 2006, Dabney-Smith & Cline, 2009, Mori et al., 2001, Pal et al., 2013). However, it is unclear what component of the cpTAT serves as the “motor” for the translocation, if any. The pulling mechanism hypothesizes that the Hcf106 and Tha4 hydrophilic domains seal the membrane to prevent proton leakage through the PMF-energized thylakoid membrane. Moreover, it has been suggested that Tha4 assembly, together with Hcf106 present in the receptor complex, is involved in the punctual weakening of the thylakoid membrane (Brüser & Sanders, 2003, Cline, 2015). In that model, Brüser and Sanders (2003) suggested that (cp)TatC could be the potential component of the (cp)Tat system that can provide a motive force to the pulling mechanism. It has been confirmed that (cp)TatC drives the substrate through the translocon (Frobel et al., 2012); our study results are consistent with that finding. The cysteine crosslinking data revealed that N-region of the N-terminal extension of cpTatC interacts with the pre-precursor mature domain (mature domain of the tOE17, which possesses signal peptide) during cpTAT translocation. According to the findings of this study, we can infer that the N-terminal extension of the cpTatC protein has a dual role: it stabilizes the signal peptide of the precursor protein and generates force to move the cargo. Firstly, we predict that the amino-proximal binding of cpTatC with the precursor mature domain stabilizes binding of the precursor and orients it such that specific recognition and binding of RR consensus motif of the signal peptide to the S1 and S2 regions of cpTatC, which in turn triggers the formation of translocon (Alami et al., 2003, Frobel et al., 2012, Gérard & Cline, 2006, Ma & Cline, 2010) in the presence of the PMF. Secondly, the amino proximal extension might provide a motive force for the mature domain of the precursor to translocate through the conduit composed of Tha4 oligomerization.

The N-terminal extension of cpTatC in chloroplasts suggests it has a specialized role different from bacterial TatC systems. It may regulate protein transport in response to diurnal changes in chloroplasts. The chloroplast cpTAT machinery needs to adapt to the cyclic nature of photosynthesis. During daylight, active photosynthesis generates a proton motive force (PMF) that powers protein transport into the thylakoid lumen. In contrast, nighttime conditions necessitate a temporary mechanism to halt or regulate protein transport through the cpTAT system. In vivo studies using GFP as a model substrate have shed light on these regulatory mechanisms (Marques *et al*., 2004, Marques *et al*., 2003). Therefore, the cpTatC-specific N-terminal extension likely contributes to chloroplast regulatory processes. It may facilitate the reversible binding and storage of stromal proteins during nighttime and play a role in modulating the activity or assembly of the cpTat complex in response to diurnal changes in photosynthetic activity. Understanding these mechanisms enhances our comprehension of chloroplast biology and provides insights into potential strategies for optimizing plant productivity and resilience in varying environmental conditions.

Upon examination of data from prior studies and our present research, we have postulated that the cpTatC N-terminal extension may play a role in stabilizing and aligning the signal peptide for interaction with the S1 and S2 loops of the cpTatC in the receptor complex (Alami et al., 2003, Frobel et al., 2012, Gérard & Cline, 2006, Ma & Cline, 2013). To accurately align the signal peptide, the N-terminal extension potentially engages initially with the early mature domain of the precursor. Further inquiry is warranted to validate whether the N-terminal extension indeed binds with the precursor protein before the S1 and S2 loops recognize the signal peptide’s RR motif. The amino-terminal extension of cpTatC is postulated to function as the stromal-proximal soluble domain, indicating its potential role as a motive force for cargo translocation. It is likened to a mobile hand, exerting pressure that propels the cargo through the translocation conduit. In summary, this study’s crosslinking data reveal that chloroplast TatC’s unique N-terminal extension plays a significant role in the higher plant Tat mechanism. Interestingly, this data shows us that different regions of the amino-terminal extension bind with different intensities with the pre-precursor mature domain. Therefore, it is worth studying the most significant region of the N-terminal extension of the cpTatC, which binds with the precursor protein during chloroplast TAT translocation.

## 5. Conclusion

The mature domain of cpTatC exhibits an N-terminal extension distinct from that of E. coli and algae TatC. The function of this extension in the chloroplast Tat system remains elusive. Our investigation employed a crosslinking method to delineate the interactions between the mature domain of the precursor (tOE17A137C) and the N-terminal extension of cpTatC, uncovering specific cysteine residue interactions. The crosslinking data indicate that diverse regions of the N-terminal extension bind with varying intensities to the precursor, underscoring its role in stabilizing and orienting the precursor protein for efficient translocation. This interaction potentially initiates the formation of the hairpin loop of the signal peptide with the cpTatC-Hcf106 receptor complex, triggering Tha4 protein assembly and the creation of a translocation conduit. These findings imply that the N-terminal extension of cpTatC stabilizes and positions the precursor’s signal peptide to engage with the stromal domains of cpTatC, thereby facilitating protein translocation. The study lends support to the hypothesis of a membrane weakening and pulling mechanism, positing that cpTatC, as a motor component, supplies the motive force for substrate translocation through the Tat translocon. Furthermore, the distinctive N-terminal extension of cpTatC in chloroplasts may govern protein transport in response to diurnal changes, adapting to the cyclic nature of photosynthesis.

## Acknowledgement

We would like to acknowledge members of the Dabney-Smith lab for critical reading of the manuscript. This work is generously supported by NIGMS/NIH R15 GM137251 and the Ernest H. Volwiler Professorship to Carole Dabney-Smith.

